# Application of Quasimetagenomics Methods to Define Microbial Diversity and Subtype *Listeria monocytogenes* in Dairy and Seafood Production Facilities

**DOI:** 10.1101/2022.11.07.515551

**Authors:** Brandon Kocurek, Padmini Ramachandran, Christopher J. Grim, Paul Morin, Laura Howard, Andrea Ottesen, Ruth Timme, Susan R. Leonard, Hugh Rand, Errol Strain, Daniel Tadesse, James B. Pettengill, David W. Lacher, Mark Mammel, Karen G. Jarvis

## Abstract

Microorganisms frequently colonize surfaces and equipment within food production facilities. *Listeria monocytogenes* is a ubiquitous foodborne pathogen widely distributed in food production environments and is the target of numerous control and prevention procedures. Detection of *L. monocytogenes* in a food production setting requires culture dependent methods, but the complex dynamics of bacterial interactions within these environments and their impact on pathogen detection remains largely unexplored. To address this challenge, we applied both 16S rRNA and shotgun quasimetagenomic (enriched microbiome) sequencing of swab culture enrichments from seafood and dairy production environments. Utilizing 16S rRNA amplicon sequencing, we observed variability between samples taken from different production facilities and a distinctive microbiome for each environment. With shotgun quasimetagenomic sequencing, we were able to assemble *L. monocytogenes* metagenome assembled genomes (MAGs) and compare these MAGSs to their previously sequenced whole genome sequencing (WGS) assemblies, which resulted in two polyphyletic clades (lineages I and II). Using these same datasets together with *in silico* downsampling to produce a titration series of proportional abundances of *L. monocytogenes*, we were able to begin to establish limits for *Listeria* detection and subtyping using shotgun quasimetagenomics. This study contributes to the understanding of microbial diversity within food production environments and presents insights into how many reads or relative abundance is needed in a metagenome sequencing dataset to detect, subtype, and source track at a SNP level, as well as providing an important foundation for utilizing metagenomics to mitigate unfavorable occurrences along the farm to fork continuum.

**IMPORTANCE:** In developed countries, the human diet is predominantly food commodities, which have been manufactured, processed, and stored in a food production facility. It is well known that the pathogen *Listeria monocytogenes* is frequently isolated from food production facilities and can cause serious illness to susceptible populations. Multistate outbreaks of *L. monocytogenes* over the last 10 years have been attributed to food commodities manufactured and processed in production facilities, especially those dealing with dairy products such as cheese and ice cream. A myriad of recalls due to possible *L. monocytogenes* contamination have also been issued for seafood commodities originating from production facilities. It is critical to public health that the means of growth, survival and spread of *Listeria* in food production ecosystems is investigated with developing technologies, such as 16S rRNA and quasimetagenomic sequencing, to aid in the development of effective control methods.

## INTRODUCTION

Food safety and quality are impacted by the microbial ecology within manufacturing facilities. Industrial surfaces (e.g., utensils, architecture, equipment) within food manufacturing facilities are known sources of microbial food contamination, acting as both vehicles for transmission to food commodities and environmental reservoirs for persistence (1–5). Additionally, raw materials, food ingredients and processing water are known to contribute to the microbiome of the finished product and may also introduce lot to lot variations (1–5).

Environmental sampling programs provide proactive and preventative surveillance strategies to identify contamination patterns in food production facilities. These programs aim to protect the food supply and maintain food quality by revealing pathogen presence and persistence and unsanitary conditions over time ([FDA Environmental Sampling](https://www.fda.gov/food/sampling-protect-food-supply/environmental-sampling#results)). Prior to sampling, each production facility is divided into zones according to the proximity of the sampling area to the food. Zone 1 includes food-contact surfaces such as processing equipment parts, food handling utensils, and tank and container interiors. Sampling areas in zones 2 through 4 are consecutively further from contact with food and present a lower risk to food safety. These include equipment parts and frames, floors and floor drains, and walls under or near equipment (zone 2) and forklifts and hand trucks (zone 3). Zone 4 areas are even more remote to food contact surfaces including areas of the facility such as cafeterias and locker rooms ([Control of *Listeria monocytogenes* in Ready-To-Eat Foods](https://www.fda.gov/media/102633/download)).

When collected for FDA inspection and environmental sampling assignments, environmental swab samples are immediately transported to an FDA regulatory field laboratory, and analyzed for microbial pathogens, such as *Salmonella*, *Listeria* and *Escherichia coli*. Pathogen isolates are then processed for whole genome sequencing (WGS) and uploaded into the GenomeTrakr database where they can be compared to food, clinical and environmental swab isolate genomes (6, 7).

Culture independent metagenomic sequencing technology has been transformative in filling knowledge gaps pertaining to the microbial ecology of clinical, food and environmental samples. Targeted 16S rRNA amplicon sequencing is a cost-effective workflow that has been used to evaluate the microbiotas in food, meat and beverage processing facilities, and their potential to impact food safety and quality, and to demonstrate temporal fluctuations of microbial communities in different types of processing facilities (1, 8–10). Some examples include the use of 16S rRNA amplicon sequencing to map microbial ecosystems in breweries (11), determine the bacterial diversity of floor drains in cheese production facilities (12) and examine the contribution of the microbiota of seafood processing facilities to spoilage and contamination of salmon fillets (13). There have also been studies in domestic built environments that investigated food preparation, such as a study on the microbiota of kitchen surfaces (14).

Shotgun metagenomic sequencing is more informative than targeted 16S rRNA amplicon sequencing since it can provide species level identification of bacterial communities and their gene content. And with sufficient genome coverage, strain level identification of foodborne pathogens can be achieved (15–17). Shotgun metagenomes can also be used to produce metagenome-assembled genomes (MAGs) of individual taxa within a dataset, when high levels of genome coverage are achieved (18, 19).

The often high bacterial burden, and complexity, of intrinsic food microbiomes, and limitations in short read sequencing depth preclude the use of culture-independent workflows, such as metagenomics, for regulatory purposes to detect pathogens in food and environmental samples. However, research laboratories are employing metagenomics to gain insights into the dynamics of culture dependent foodborne pathogen detection methods. Currently, quasimetagenomic approaches including culture enrichment alone, or in conjunction with techniques that shorten enrichment times such as, multiple displacement amplification (MDA) and immunomagnetic separation (IMS), have been employed to amplify pathogen genomic signatures in foods (20–24).

In this study 16S rRNA amplicon and quasimetagenomic sequencing were performed on environmental swab culture enrichments from dairy and seafood production facilities. The 16S rRNA amplicon data was used to identify differences in the microbiotas within and among the firms. SNP analyses was performed to compare *L*. *monocytogenes* MAGs from shotgun datasets to their corresponding WGS *L*. *monocytogenes* genomes from the GenomeTrakr Pathogen detection database. An *in silico* downsampling approach was then employed to reduce the percentage of sequence reads in shotgun metagenomes that varied in their proportional abundances of *L*. *monocytogenes* to test how a reduction in *L*. *monocytogenes* sequence reads would impact SNP clustering. Taken together, this study revealed microbiota differences from 12 dairy and seafood facilities and variations in abundances of *L*. *monocytogenes* subtypes and *Listeria* species that impacted SNP clustering. Additionally, we identified bacteria that co-enrich with *Listeria* in Modified *Listeria* Enrichment Broth (UVM) with antimicrobial supplements, including some that may affect food safety and quality.

## MATERIALS & METHODS

### Samples used in this study

Our sequence datasets were produced from 355 environmental swab UVM *Listeria monocytogenes* culture enrichments from seven dairy (245 samples) and five seafood (110 samples) production facilities (Table 1). After the regulatory culture work was completed, 2 mL aliquots were removed from randomly chosen culture positive and negative 48-hour UVM culture enrichments for metagenomic DNA extraction. Aliquots were centrifuged at 12,000 x g for 10 min, and bacterial pellets were stored at −20°C until DNA extraction was performed. All aliquots for metagenomic analysis were taken after the FDA regulatory assignments were completed, when the samples were no longer needed for regulatory analysis and disposition. Aliquots from earlier timepoints of the regulatory workflow, such as un-enriched microbiomes, were not available due to chain of custody policies.

**Table 1.** Environmental swab culture enrichments analyzed with 16S rRNA amplicon and shotgun metagenomic sequencing from 7 dairy and 5 seafood manufacturing facilities.

| Firm | Food Type<sup>a</sup> | 16S rRNA<sup>b</sup> | Shotgun sequenced<sup>c</sup> | % Lm Pos<sup>d</sup> | Zone 1<sup>e</sup> | Zone 2<sup>e</sup> | Zone 3<sup>e</sup> |
| --- | --- | --- | --- | --- | --- | --- | --- |
| AC | D | 12 | 1 | 0 | NA | NA | NA |
| AM | D | 68 | 7 | 31 | 18 | 21 | 29 |
| G | D | 45 | 7 | 44 | NA | 6 | 21 |
| I | D | 54 | 7 | 9 | 8 | 15 | 31 |
| J | D | 39 | 3 | 36 | 13 | 26 | 0 |
| O | D | 18 | 1 | 22 | 2 | 1 | 15 |
| V | D | 9 | 1 | 0 | NA | NA | NA |
| A | S | 32 | 7 | 13 | 27 | 2 | 3 |
| AS | S | 12 | 1 | 33 | NA | NA | NA |
| F | S | 24 | 6 | 67 | 2 | 7 | 15 |
| L | S | 29 | 9 | 10 | 13 | 5 | 11 |
| W | S | 13 | 2 | 31 | 0 | 5 | 8 |

<sup>a</sup>Dairy (D) or Seafood (S)
<sup>b</sup>Number of samples analyzed with 16S rRNA amplicon sequencing
<sup>c</sup>Number of samples analyzed with shotgun metagenomic sequencing
<sup>d</sup>Percent of 16S rRNA samples culture positive for *Listeria monocytogenes*
<sup>e</sup>Number of samples collected from each environmental monitoring zone; no zone 4 samples were collected
NA - zone information not available

Since FDA procedures for culture-positive samples require whole genome sequencing of pathogen isolates followed by entry of pathogen genomes into the GenomeTrakr database (NCBI PRJNA215355), the culture results of the dairy and seafood environmental swab enrichments were known prior to metagenomic sequencing (Table 1 and Table 2). Of the 355 samples sequenced in this study, 65 diary and 33 seafood samples were culture-positive for *Listeria monocytogenes* (Table 2). Some UVM culture enrichments were also culture-positive for *L*. *innocua* (21 samples), *L*. *welshimeri* (1 sample), *L*. *seeligeri* (1 sample), and *L*. *ivanovii* (1 sample); these isolates were not sequenced or uploaded into the GenomeTrakr database (Table 2).

**Table 2.**
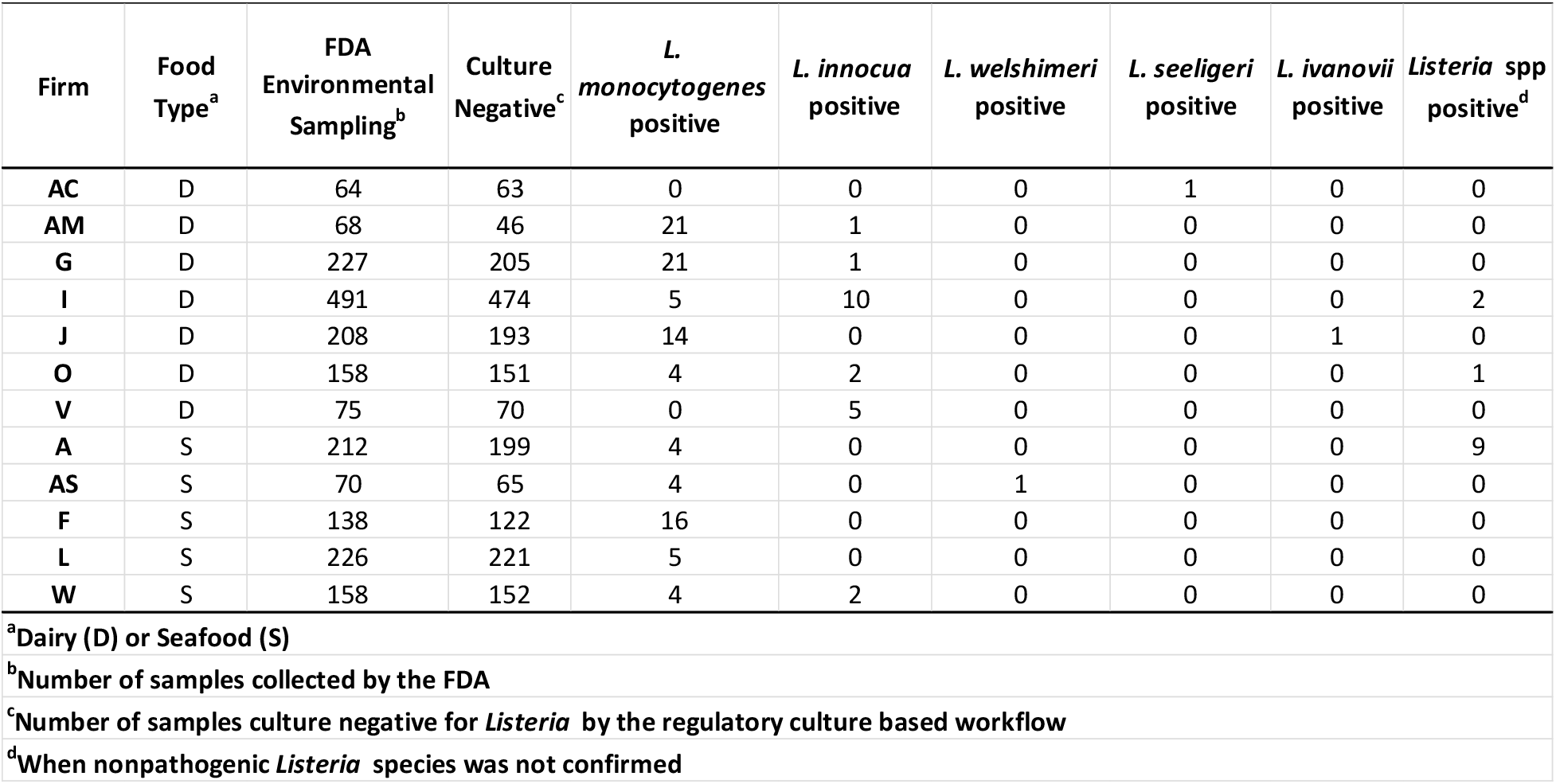
Culture enrichment results for environmental swab samples from 7 dairy and 5 seafood manufacturing facilities.

### Preparation of Genomic DNA

Genomic DNA was extracted from bacterial pellets using either the Qiagen DNeasy Blood and Tissue Kit (Qiagen, Germantown, MD, United States) with modifications (92 bacterial pellets), or the ZymoBIOMICS® DNA Miniprep Kit (Zymo, Irvine, CA, United States) (263 bacterial pellets). For samples processed with the DNeasy kit, the Gram-positive protocol was modified to incorporate a pre-lysis step. Bacterial pellets were resuspended with 170 μL of enzymatic lysis solution (20 mM Tris-HCl (pH 8.0), 2 mM Sodium EDTA, 1.2% Triton X-100, filter sterilized + 100mg/mL Lysozyme), incubated for 2 hours at 37°C and then boiled at 95°C for 15 minutes. After cooling for 5 minutes, 10 μL of 10% SDS was added.

DNA extraction with the ZymoBIOMICS® DNA Miniprep Kit was performed according to the manufacturer’s specifications, using mechanical lysis for 40 continuous minutes with a Vortex Genie 2 (Scientific Industries, Bohemia, NY, United States) and a horizontal adaptor. All samples were stored at −20°C until further processing. Library preparation for 16S rRNA amplicon and shotgun metagenomic sequencing used the same metagenomic DNA.

### 16S rRNA amplicon sequencing method

A two-step amplification protocol similar to the published Illumina protocol ([Illumina 16S Metagenomic Sequencing Library Preparation](https://support.illumina.com/documents/documentation/chemistry_documentation/16s/16s-metagenomic-library-prep-guide-15044223-b.pdf)), was employed for 16S rRNA amplicon library preparation. The first amplification step targeted the V1 to V3 hypervariable regions of the 16S rRNA gene with the forward primer 26F (5’ TCG TCG GCA GCG TCA GAT GTG TAT AAG AGA CAG DAG AGT TTG ATC MTG GCT CAG 3’) and reverse primer 534R (5’ GTC TCG TGG GCT CGG AGA TGT GTA TAA GAG ACA GTM TTA CCG CGG CNG CTG GCA C 3’) (Integrated DNA Technologies). PCR products were purified with Agencourt AMPure XP beads (Beckman Coulter, Brea, CA, United States) at 1.25× sample volume and quantified using the Qubit 2.0 fluorometer (Invitrogen, Carlsbad, CA, United States).

The second round of PCR was performed with Illumina Nextera XT indexing primers (Illumina Inc., San Diego, CA) to generate amplicons compatible with Illumina MiSeq sequencing chemistry. Library pools (8pM) were spiked with 10% PhiX Control v3 (8pM) (Illumina Inc., San Diego, CA, USA). The 355 libraries were sequenced with additional libraries from a larger study, in 12 MiSeq runs, multiplexed at 72-96 samples per run in paired-end mode with 325 × 275 cycles using the MiSeq Reagent v3 600-cycle kit (Illumina Inc., San Diego, CA).

### Processing and taxonomic assignment of 16S rRNA gene amplicons

Preprocessing and classification of reads were done as follows. First, we used FLASH (25) to merge paired-end reads keeping read 1 when the reads could not be merged (either due to insufficient overlapping basepairs or poor quality within the overlapping region). We then used the program USEARCH (26) to perform subsequent quality filtering to prepare the 16S rRNA sequence reads for taxonomic classification. This included using the fastx_truncate option to remove the primers, fastq_filter to filter based on a minimum fragment length of 150bp and ee (expected error) value of 1.0 quality, fastx_uniques to get unique sequences per sample, and unoise3 to check for chimeras. The resulting sequences were then classified using MAPSeq with the default database created using NCBI GenBank and RefSeq sequences annotated as ribosomal RNA with 16S or 18S in the annotation (27).

### Shotgun metagenomic sequencing method

Shotgun metagenomic sequencing libraries were prepared using either the Illumina Nextera XT Library Preparation Kit (39 samples) or the Illumina DNA Prep library Kit (Illumina Inc., San Diego, CA) (13 samples). Libraries were sequenced in 8 MiSeq runs multiplexed at 6-8 samples per sequencing run using MiSeq v3 600-cycle chemistry with 2 × 300 cycles.

### k-mer identification of shotgun metagenomes

Determination of the bacterial composition in shotgun metagenomes from environmental swab culture enrichments was conducted using custom C++ programs developed to compile a k-mer signature database containing multiple unique 30 bp sequences per species, and then each read in the metagenomic fastq file was identified using the 30 bp k-mer probes. For each bacterial species or subspecies, each non-duplicated 30-mer from a reference whole genome sequence was placed into a database. The k-mers not found in at least 2/3 of a set of additional genome sequences of the same species, and k-mers found in genomes of other species were removed. The resulting k-mer database used in this work contains 5900 target entries, each consisting of approximately 40,000 (range 44 to 80,000) unique k-mers. The database includes 1100 different bacterial genera and 3500 species. Normalization is performed to correct for bias due to differing number of k-mers used per database entry and the results are tabulated as percent of identified reads (contribution to the microbial population of identified species) for each database entry (github.com/mmammel8/kmer_id) (17).

### Assembly of shotgun reads

Sequence reads from isolated colonies of *Listeria* were assembled *de novo* into contigs using SPAdes (Center for Algorithmic Biotechnology, St. Petersburg State University, St. Petersburg, Russia) genome assembler version 3.13.0 with default parameters. Reads from shotgun metagenomic sequencing were assembled into contigs using SPAdes assembler with the --meta option. *Listeria* contigs were identified by nucleotide BLAST (megablast) of the assembled contigs against reference genomes of *Listeria*. Contigs matching any of ten *L. monocytogenes* reference genomes including serotypes 1/2a, 1/2b, 1/2c, 3a, 4a, and 4b, at a nucleotide identity of at least 88% were retained for further analysis (see Table S1 in the supplemental material).

### SNP alignment of *Listeria* WGS and metagenomic assemblies

To determine the core gene set for SNP alignment, 2819 annotated genes of *L. monocytogenes* str. 4b F2365 (GenBank accession AE017262.2) were used as references for BLAST analyses against 385 representative whole genome sequences of *Listeria* species *L. monocytogenes*, *L. innocua*, *L. ivanovii*, *L. marthii*, *L. seeligeri*, and *L. welshimeri*. A gene was excluded if more than one match occurred in a genome or if the gene was absent in more than one of the representative genomes at a 75% sequence identity cutoff. The resulting core backbone contained 1650 genes covering 1.5 Mbp. Next, the 1650 genes were aligned by BLAST analysis against the WGS, and metagenome assemblies generated from isolates and samples in this study, respectively. Each assembly’s matching sequence to the reference query was added to an alignment for that gene. A C++ program and a Python program (which are available on GitHub, https://github.com/mmammel8/core_genes) were used to scan each alignment and produce a consensus sequence.

### Phylogenetic analysis

Concatenated backbone SNPs were imported into MEGA 3.1 for phylogenetic analysis (28). Neighbor joining trees were constructed using the p distance model of nucleotide substitution and pairwise deletion option. The SNP distances are estimates based on the proportion of differences for the pairwise comparisons between samples. The proportions were multiplied by the number of polymorphic backbone sites within each dataset (13,195 for *L. monocytogenes* lineage I and 25,556 for *L. monocytogenes* lineage II) and then rounded to the nearest integer to get the estimated number of differences. This is necessary because of gaps (missing data) in the backbone gene alignments from the assemblies.

### *In silico* analysis of *Listeria* in culture positive shotgun metagenomes

SNP analyses of the culture positive metagenomic assemblies provided insights into the sequencing depth required to group *L*. *monocytogenes* metagenome-assembled genomes (MAGs) with their corresponding WGS genomes. Our next goal was to investigate how varying the number of *L*. *monocytogenes* sequence reads in MAG data sets impacts SNP calling. We employed an *in silico* downsampling process to reduce the total number of sequence reads, in three lineage I culture positive environmental swab enrichments that varied in proportional abundances of *L*. *monocytogenes*. The first metagenomic fastq file S105 from dairy firm I, with 5,196,513 sequence reads, (86% proportional abundance of lineage I *L*. *monocytogenes*) was reduced by downsampling from the original set at a probability of 0.5% to 8% to generate 7 additional fastq files. For all these individual fastq files, the contigs were assembled using SPAdes. The assembled contigs were added to the core gene alignments to determine the SNPs and added to a SNP tree. The same method was followed for MAG S170 from dairy firm G and MAG S064 from dairy firm I with 44% and 0.106% abundances of *L*. *monocytogenes*, respectively. These samples had reads selected at percent ranges that would provide sufficient numbers of *Listeria* reads for assembly (2-16% and 86-66%, respectively).

#### Statistics

Following OTU clustering and taxonomic assignments, alpha diversity (Observed OTUs) and beta diversity were calculated and assessed using the following R packages: Bioconductor, metagenomeSeq, vegan, phyloseq, and ggplot2. Significance tests were conducted using an ANOVA, ADONIS and pairwise ADONIS tests. A value of *p* less than 0.05 were considered statistically significant. Beta-diversity was examined using Bray-Curtis dissimilarity matrix. Principal component analysis for Aitchison compositions were used when appropriate. Alpha diversity metrics were measured using Shannon diversity index with significance determined by the non-parametric Kruskal Wallis tests. False Discovery Rate (FDR) correction was used to correct for multiple hypothesis testing. All statistical analyses were conducted in R Studio (version 1.1.456, using R version 3.6.3). The visualizations were created using ggplot2 unless otherwise mentioned in the figures.

### Database Submission

16S rRNA amplicon and shotgun metagenomic sequencing data from this study has been submitted to the FDA MetagenomeTrakr Project (NCBI Accession PRJNA 540844). The data from this package was submitted using MIxS food-food production metadata package.

## RESULTS

### 16S rRNA Amplicon Sequencing

16S rRNA amplicon sequencing of environmental swab culture enrichments from the 12 seafood and dairy manufacturing firms yielded microbiomes that varied in alpha diversity (Fig. 1). However, none of the variation was significant, likely due to the culture bias introduced during enrichment. The enrichment process included antimicrobials which favored the growth of gram-positive organisms, including *Listeria* species, and had an antagonistic effect against the proliferation of gram-negative bacteria. Additionally, no significant differences were observed in the Shannon diversity of *L*. *monocytogenes* culture positive (n=95) and culture negative (n=260) enrichment microbiomes (p = 0.75) (Fig. 2); however, observationally, the culture *L. monocytogenes* positive enrichment microbiomes appeared to have higher diversity. Principal Component Analysis (PCA) of the Bray-Curtis distances, did not reveal significant differences (adjusted P = 1) in beta diversity among the 245 dairy and 110 seafood culture enrichments from the 12 firms (Fig. 3), nor those that were culture positive or negative for *L. monocytogenes* (see Fig. S1 in the supplemental material).

**Figure 1.**
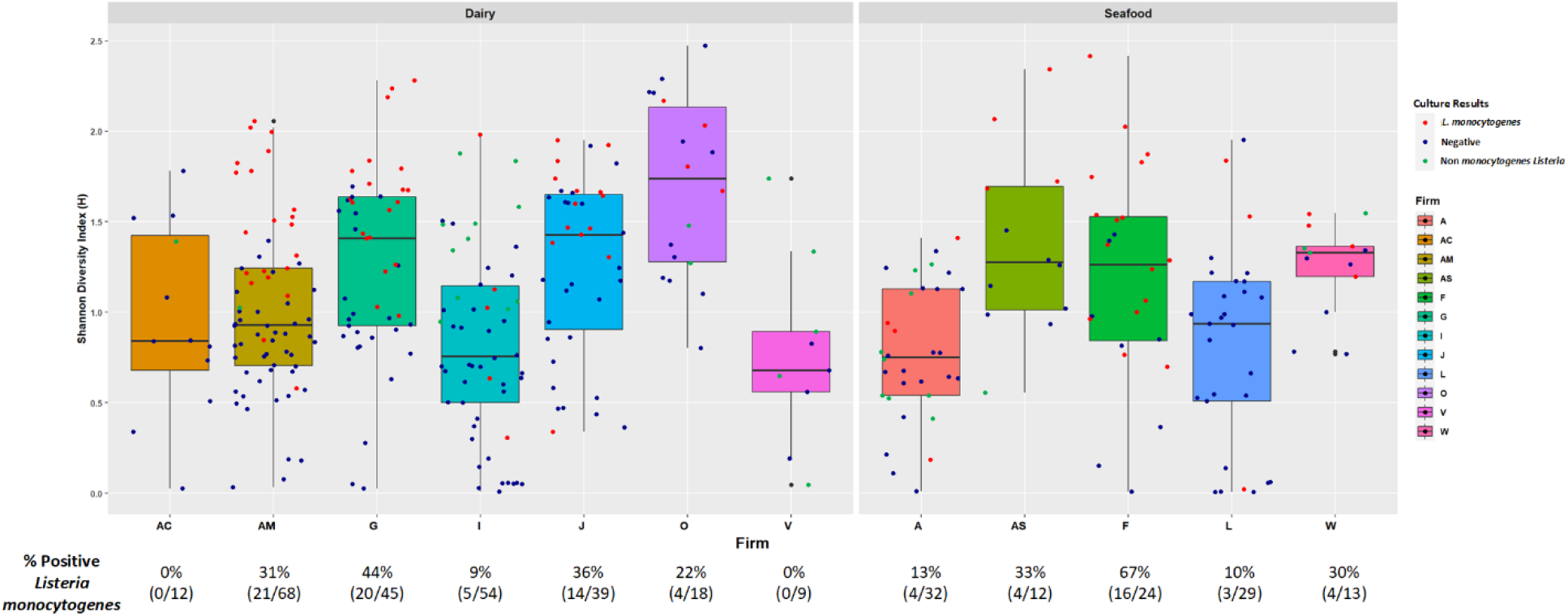
Shannon diversity of enrichments for each sample collected from the seven dairy firms (left panel) and 5 seafood firms (right panel). Color of the dot indicates the regulatory culture result (blue = *Listeria* culture negative, red = *Listeria monocytogenes* culture positive, green = culture positive for non-monocytogenes *Listeria*). Percentage of *L. monocytogenes* culture positive samples are indicated at the bottom.

**Figure 2.**
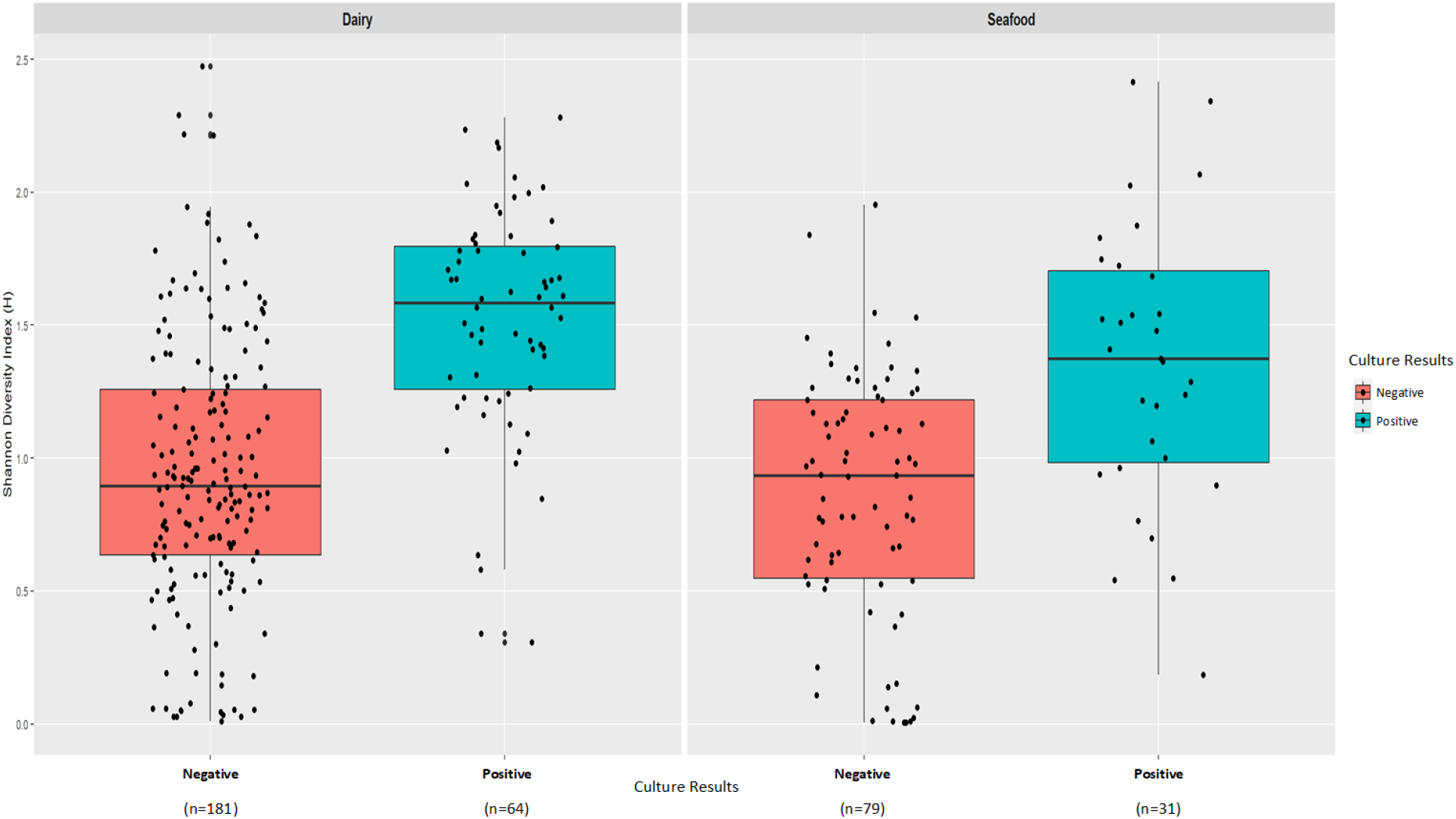
Shannon diversity of enrichments of the 355 samples via box plots faceted by the manufacturing facility type Dairy (left panel) and Seafood (right panel) and culture results for *L. monocytogenes* (orange = culture positive, blue = culture negative); n = number of swab enrichments.

**Figure 3.**
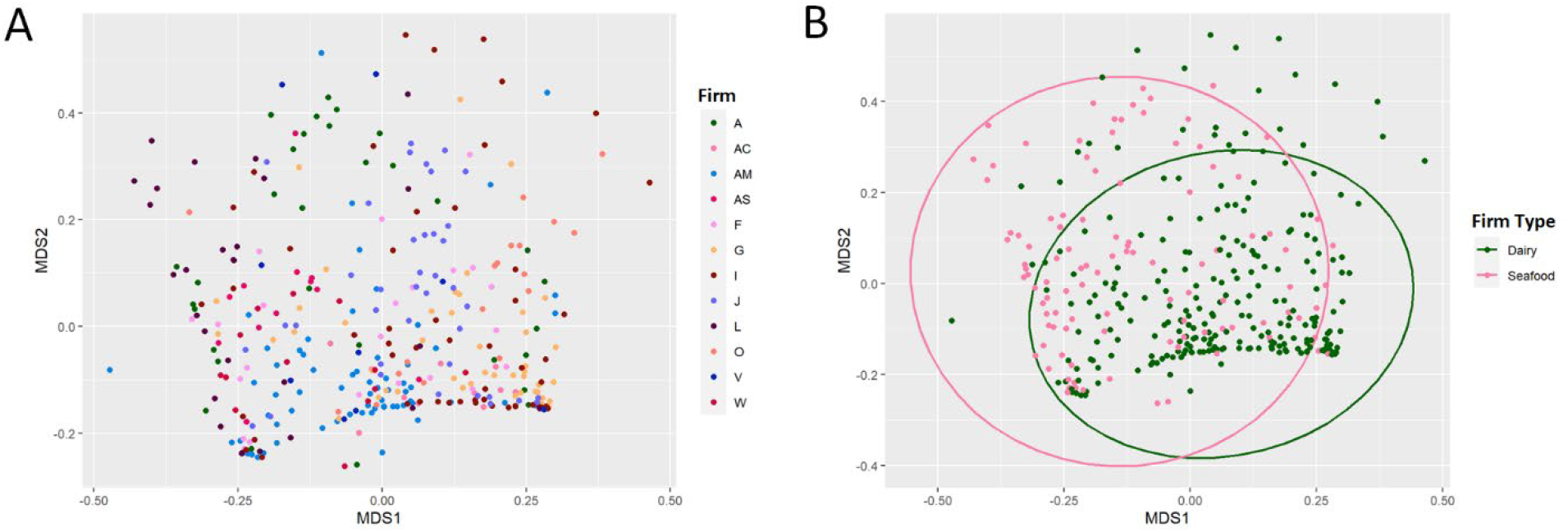
Principal Component Analysis (PCA) of Bray Curtis dissimilarity distances between the 355 UVM culture enrichments. (A) Beta-diversity between samples collected from the 12 food manufacturing firms (AC, AM, G, I, J, O and V = dairy firms; A, AS, F, L and W = seafood firms). (B) Beta-diversity of samples collected from dairy manufacturing firms (n=245) and seafood manufacturing firms (n=110).

The potential biases in microbiome diversity introduced by UVM culture enrichment prompted us to test a second beta diversity metric, Aitchison PCA, to account for the possibility of decreased diversity in the enriched microbiomes. The Aitchison PCA metric showed that beta-diversity was driven by a dominance of *Lactococcus*, *Enterococcus* and *Pseudomonas* for the culture negative samples and *Listeria*, and *Vagococcus* for the culture positive samples. All of these taxa are within the Gram-positive Firmicutes phyla except *Pseudomonas*.

The UVM culture enrichment of the environmental swabs from the 12 dairy and seafood firms resulted in bacterial communities dominated by either Firmicutes or Proteobacteria followed by either Actinobacteria, Fusobacteria or Bacteroidetes (Fig. 4; see Table S2 in the supplemental material). Variations in proportional abundances of Actinobacteria, Fusobacteria or Bacteroidetes phyla were observed in some firms. For example, Fusobacteria 16S rRNA sequence reads were identified in dairy firms O, AM, G and I, and seafood firm F at proportional abundances ranging from 17% to 46% and 12 to 37%, respectively, all classified as *Fusobacterium* (Fig. 4, see Table S2 in the supplemental material).

**Figure 4.**
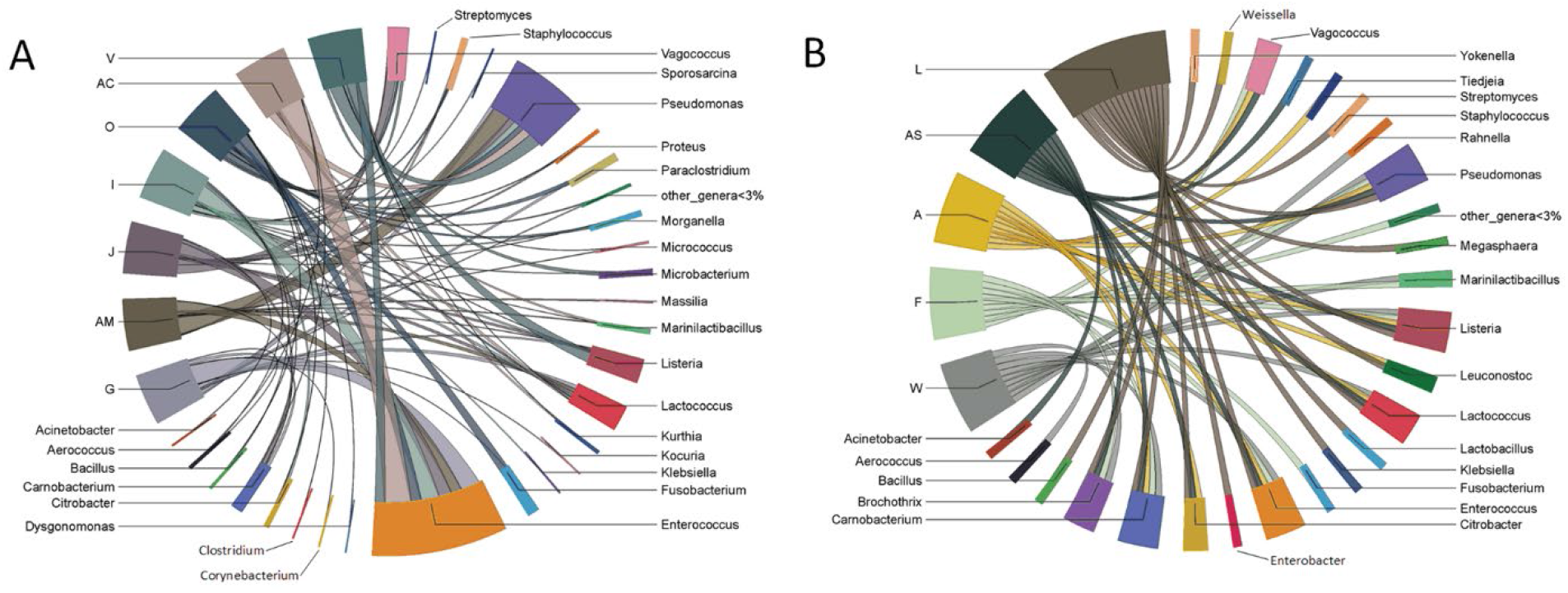
Chord diagrams of shared taxa found in environmental swab culture enrichments taken from 7 dairy (A) and 5 seafood (B) facilities. All genera less than 3% relative abundance after 16S rRNA amplicon MAPseq analysis have been incorporated into “other_genera<3%”.

Actinobacteria were observed in 64% (226/355) of all swab culture enrichments. The dairy firms had more samples with Actinobacteria at higher proportional abundances (60% to 96% in some samples) than those from seafood firms (see Table S2 in the supplemental material). For example, Actinobacteria was present in 100% of diary firm G samples, which were collected from the firm on three sampling days during 2018 and 2019. The predominant Actinobacteria taxa for each collection day were *Kocuria*, on 04/10/2018, *Rothia* on 04/08/2019, and *Corynebacterium* on 04/08/2019 (see Table S2 in the supplemental material).

Proportional abundances of Bacteroidetes, namely *Dysgonomonas*, were low (8% to ≤1%) in both the dairy and seafood firm enrichments (Fig. 4; see Table S2 in the supplemental material). The Firmicutes population in dairy firm culture enrichments were dominated by *Enterococcus*, *Lactococcus*, and *Vagococcus*, which impacted the Aitchison PCA beta diversity scores, although *Carnobacterium* predominated in enrichments from some swab samples (Fig. 4; see Table S2 in the supplemental material).

*Pseudomonas* was the dominant taxa in the Proteobacteria phylum in both the dairy and seafood firms (Fig. 4; see Table S2 in the supplemental material). Differential abundance analyses between the dairy and seafood firms revealed that *Brochothrix*, *Carnobacterium*, *Listeria*, *Leuconostoc*, *Yokenella*, and *Tiedjeia* were significantly higher in dairy firms (p <0.05). Furthermore, *Bacillus*, *Aerococcus*, *Massilia*, *Staphylococcus*, *Proteus*, *Fusobacterium*, *Morganella*, *Paraclostridium*, and *Enterococcus* abundances were significantly lower (p < 0.05) in dairy firms compared to seafood firms.

### Bacterial Fingerprints of Taxa in Dairy Firm AM

To determine the potential of employing quasimetagenomics to map the microbiota in food manufacturing firms, a more thorough analysis of culture enrichments from dairy firm AM was performed. This firm was chosen for its’ comprehensive metadata, which was used to create a representative floorplan enabling the identification of the environmental swab enrichment microbiomes in different geographic regions and zones within the firm (Fig. 5). Dairy firm AM produces various cheeses and cheese products, and 68 environmental swabs were collected from either the cleaning (n=13) or processing (n=55) rooms and grouped by proximity into 10 areas (A – J) (Table 1; Fig. 5). *L*. *monocytogenes* was cultured by regulatory analysts from swabs collected in eight of the ten areas (Table 1; Fig. 5). Both *L. innocua* and *L*. *monocytogenes* were cultured from one swab collected in area F, and all swabs from area G were culture negative for *Listeria*.

**Figure 5.**
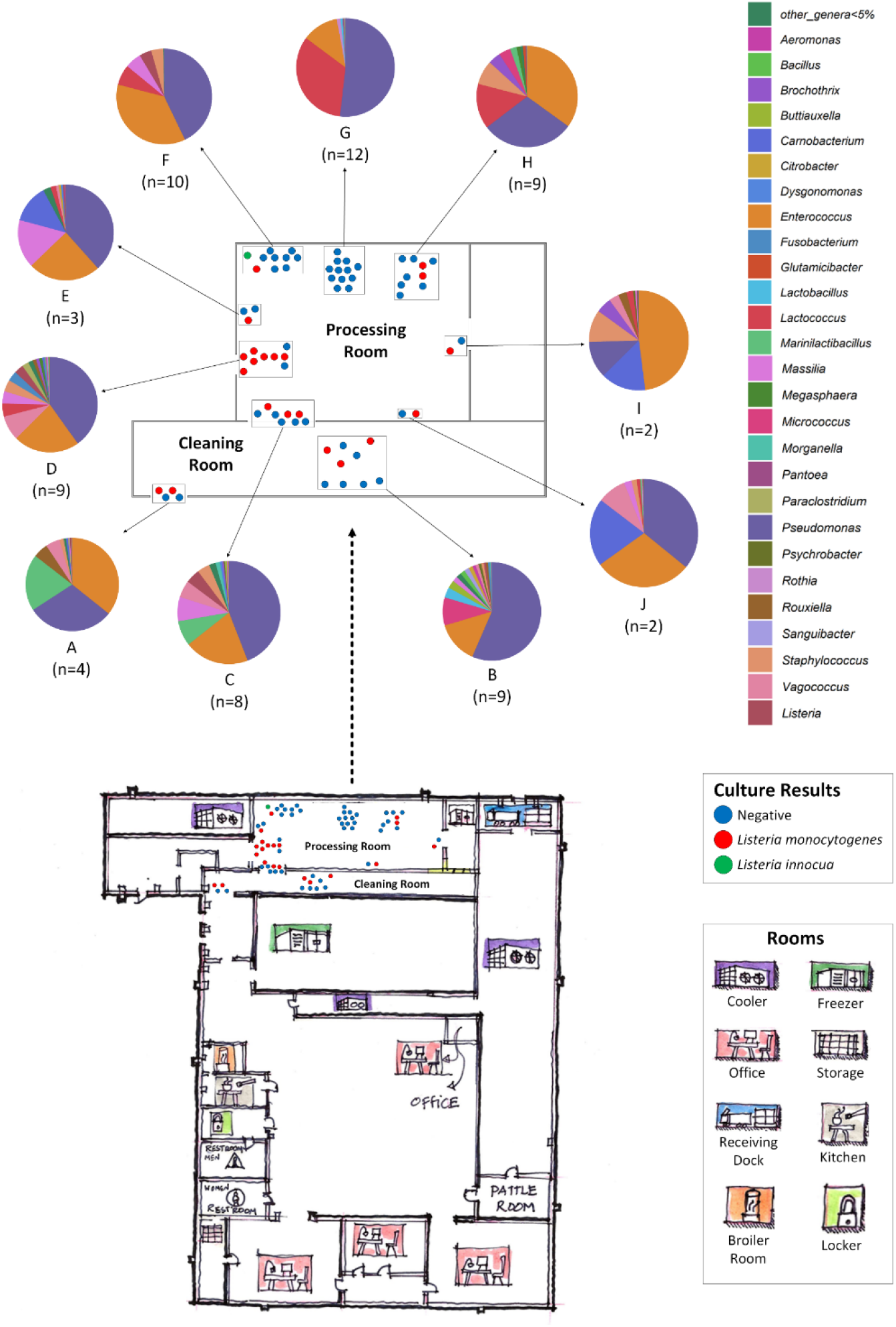
Hypothetical floorplan of dairy production facility FirmAM. Room legend indicates separate rooms of the facility where a specific step in food production occurs. The relative areas sampled (dots) are marked and color coded to reflect the regulatory culture result (blue = negative, red = *Listeria monocytogenes* positive, green = *Listeria innocua* positive). The swab collection sites (n=68) were grouped into 10 sampling areas (A, B, C, D, E, F, G, H, I, J) within the cleaning and processing rooms. This floorplan is not drawn to scale.

The 16S rRNA microbiota of the culture enrichments from the 10 sampling areas were dominated by either *Pseudomonas* (13-56% proportional abundances) or *Enterococcus* (11-47% proportional abundances) (Fig. 5). However, taxa present at ≤12% relative abundances in the cleaning and processing rooms differed. The processing room harbored *Lactococcus* (12%), *Massilia* (4%), and *Staphylococcus* (4%), while the cleaning room areas harbored *Micrococcus* (7%), *Marinilactibacillus* (6%), and *Lactobacillus* (2%) (Fig. 5). Samples from area B of the cleaning room were the most diverse with 23 taxa observed at proportional abundances ≥1% in at least one sample. Area G in the processing room, with the highest number of samples (n=12), harbored the least diverse bacterial communities in enriched samples with only 10 taxa at proportional abundances ≥1% in at least one sample and this area had the highest average proportional abundances of *Lactococcus* (33%); all 12 enrichments were culture negative for *L*. *monocytogenes*. Additionally, *Carnobacterium* proportional abundances were ≥12% in the processing room areas E, I & J. *Vagococcus* was observed in all areas except G and H in the processing room, at proportional abundances ranging from 1% to 9% in at least one sample (Fig. 5).

PCA plots of the Bray-Curtis dissimilarity distances among the microbiotas from firm AM culture enrichments revealed that the beta diversities of area G, in the processing room, significantly differed from swabs collected in areas C (p = 0.045) and D (p = 0.045) also from the processing room, as well as areas A (p = 0.045) and B (p = 0.045) from the cleaning room (Fig. 6). These differences were likely due to the high proportional abundances of *Lactococcus* and low diversity of bacteria in area G (Fig. 5). The swabs collected from area G were collected from zone 1 food contact surfaces such as utensils and commonly touched items such as mixing equipment and the outer surfaces of vessels used for processing designated as zone 2. Since firm AM manufactures and processes cheese it is not surprising that the highest abundances of *Lactococcus* were observed in the processing areas (Fig. 5).

**Figure 6.**
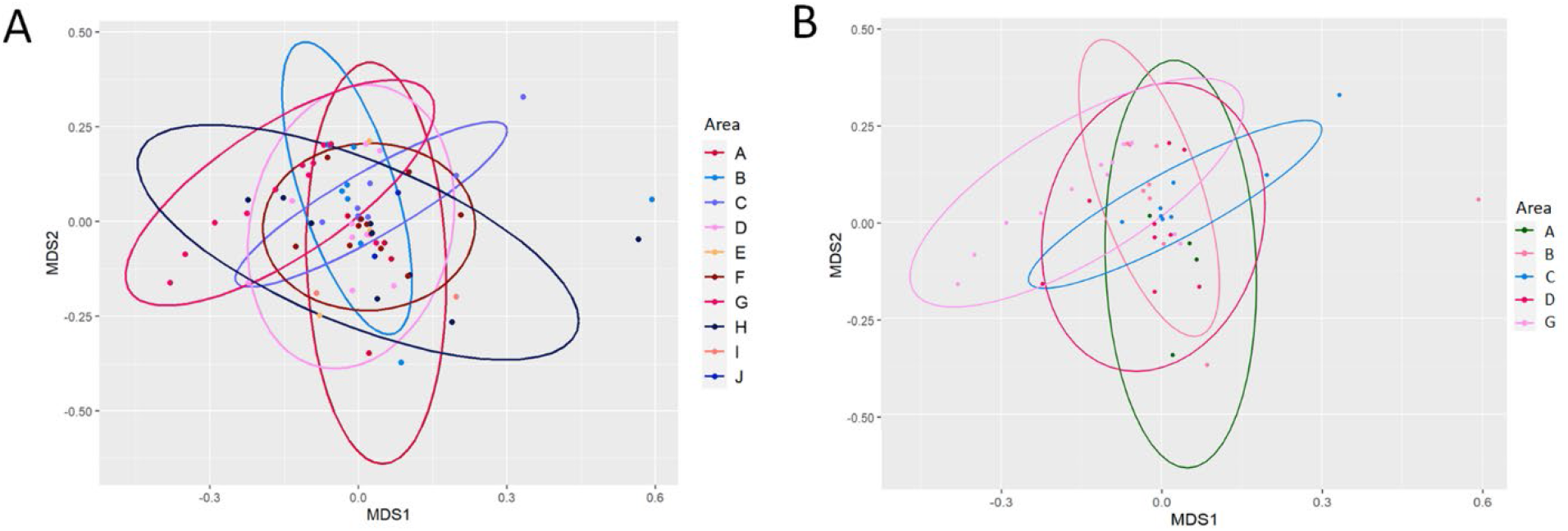
Principal Component Analysis (PCA) of Bray Curtis dissimilarity distances from culture enrichments from dairy firm AM cleaning and processing rooms. (A) Beta-diversity between all samples taken from the 10 areas. (B) Areas A (n=4), B (n=9), C (n=8) and G (n=12) had significant differences in beta diversities when compared to the other 6 areas (D, E, F, H, I, J).

### Shotgun Metagenomic Analyses

Shotgun metagenomic sequencing of 52 environmental swab culture enrichments provided species level identification of the microbiota in a subset of samples from the dairy and seafood firms. *Enterococcus* species were more prevalent in dairy firm culture enrichments, and predominated by *E*. *casseliflavus* (see Table S3 in the supplemental material). K-mer analyses also identified *E*. *durans*, *E*. *faecalis*, *E*. *faecium*, and *E*. *mundtii*. *P*. *putida* and *P*. *fluorescens* were the most prevalent *Pseudomonas* species in dairy culture enrichments (Sup Table S2). *Carnobacterium maltaromaticum* was present in a few dairy firms at proportional abundances ≤ 20% and *Brochothrix thermosphacta* was identified in dairy firm AM microbiomes at proportional abundances ≤ 4% (see Table S3 in the supplemental material). K-mer analyses of UVM swab enrichments from seafood firms also identified multiple species of *Enterococcus* including *E*. *casseliflavus*, *E*. *faecalis* and *E*. *aquimarinus*, and *P*. *fluorescens* was the most predominant *Pseudomonas* species (see Table S3 in the supplemental material). Additionally, the prevalence and proportional abundances of *C*. *maltaromaticum* and *B*. *thermosphacta* were higher in seafood firm culture enrichments (see Table S3 in the supplemental material).

### SNP Phylogeny of genomic and metagenomic *Listeria monocytogenes* assemblies

To investigate the concordance of *L. monocytogenes* SNP phylogenies from culture positive metagenomes and corresponding WGS isolates, we selected and analyzed 32 dairy and seafood firm enrichments. Since the isolate genomes from each sample were available in the GenomeTrakr database we were able to simultaneously perform SNP analysis of the WGS and corresponding *L*. *monocytogenes* metagenomic assemblies (MAGs).

SNP analysis of the *L. monocytogenes* MAGs from 28 of the 32 dairy and seafood swab samples and their corresponding WGS isolate genomes resolved two polyphyletic clades consisting of *L*. *monocytogenes* lineages I and II with 13,195 and 25,556 SNP sites, respectively. The remaining four MAG samples did not produce adequate *L*. *monocytogenes* genome coverage to be placed in either SNP clade, although their corresponding WGS assemblies were placed in the tree (Fig. 7). Both lineage I and II *L*. *monocytogenes* were cultured from seafood firms F and L, and dairy firm G (Fig. 7). Sample S064 from dairy firm I was culture positive for lineage I *L*. *monocytogenes* and *L*. *innocua* (Fig. 7). Additionally, the majority of lineage I assemblies are from swabs collected in dairy firms (12/13), while the lineage II clade is mostly from seafood firms (12/15) (Fig. 7). Refer to Supplemental Table S4 in the supplemental material for bacterial kmer analysis results and associated metadata of MAGs.

**Figure 7.**
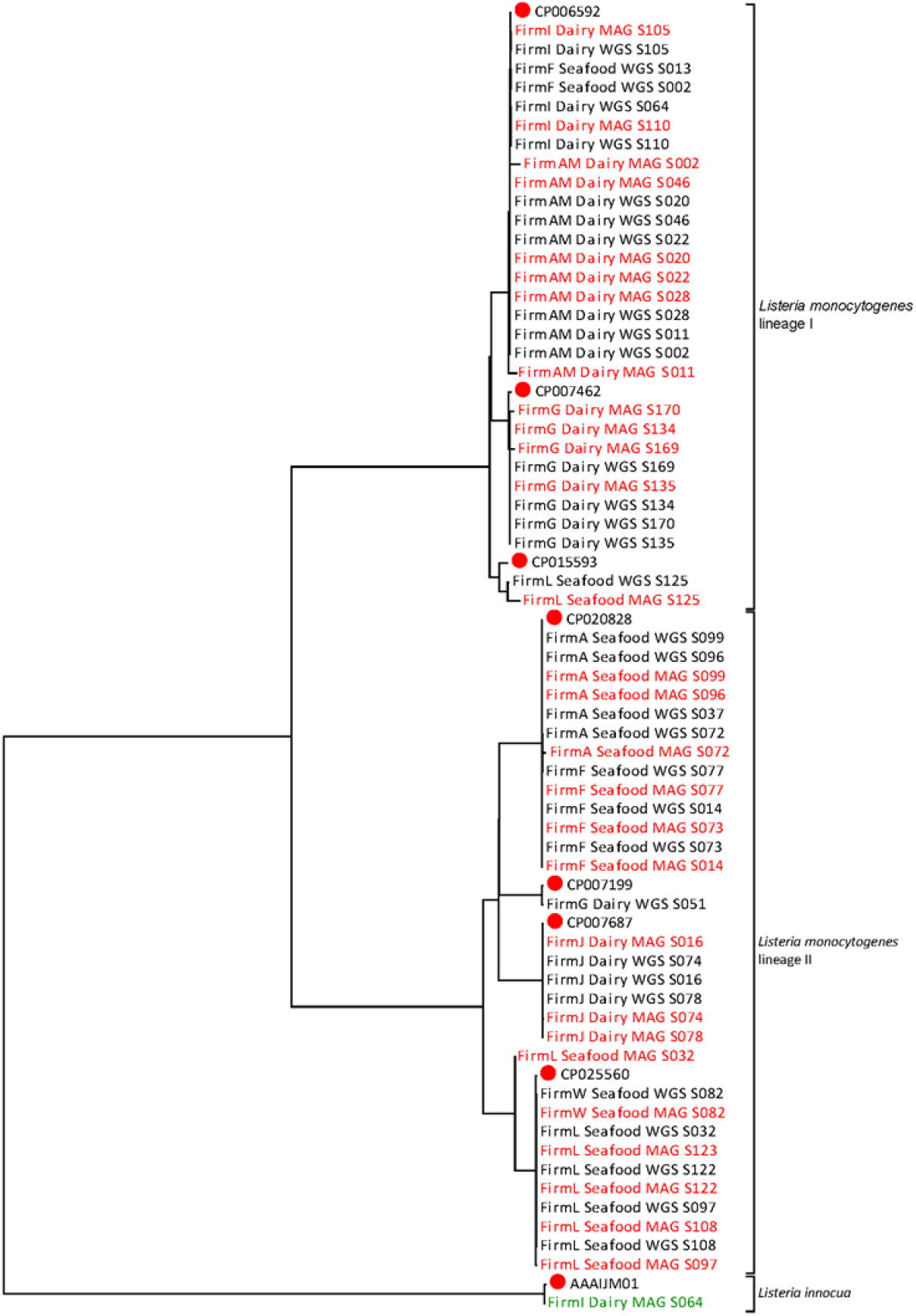
Phylogeny of *Listeria* pure isolates (WGS) and metagenomic assembled genomes (MAG) determined by single nucleotide polymorphism (SNP) analysis. WGS and MAG are designated by name and color (*Listeria monocytogenes* MAG = red, *Listeria monocytogenes* WGS = black, *Listeria innocua* MAG = green). Samples were named according to the blinded firm name followed by the designation of dairy or seafood manufacturer, whether the sample was sequencing using shotgun metagenomics (MAG) or pure isolate whole genome sequencing (WGS) and the environmental swab number. Samples that resolved to *Listeria monocytogenes* lineage I, lineage II and *Listeria innocua* are included in the tree.

The lineage I clade of the SNP tree has three subclades consisting of ten MAGs that group with their corresponding *L*. *monocytogenes* WGS genome assemblies (Fig. 7; see Fig. S2A in the supplemental material). The CP006592 subclade includes WGS and MAG assemblies from dairy firms AM and I, and seafood firm F. The WGS assemblies of dairy firm AM samples S011 and S002 grouped within the CP006592 subclade, but the corresponding MAG assemblies both grouped outside of the clade, likely due to low SNP representation (Fig. 7, see Fig S2A and Table S4 in the supplemental material). The remaining dairy firm AM WGS and MAG assemblies in the CP006592 subclade are from samples S020, S022, S028 and S046 (Fig. 5, Fig. 7, see Fig. S2A in the supplemental material).

Three samples in the CP006592 subclade were collected from dairy firm I. Two of them, produced MAG and WGS *L*. *monocytogenes* assembles that grouped together in this subclade (Fig. 7, see Fig. S2A in the supplemental material). The WGS assembly of S064 grouped within the CP006592 subclade, but MAG S064 did not, likely due to low genome coverage (Fig. 7, see Fig. S2A, Table S3 and Table S4 in the supplemental material). Nevertheless, sample S064 was also culture positive for *L*. *innocua*, and the MAG contained sufficient genome coverage to group with the *L*. *innocua* reference strain AAAIJM01 (Fig. 7). The final two samples in the CP006592 subclade, S002 and S013, did not have sufficient genome coverage for MAG assemblies but the WGS isolate genomes were placed in this subclade (Fig. 7, see Fig. S2A and Table S4 in the supplemental material).

A second lineage I SNP subclade comprised of WGS and MAG assemblies from dairy firm G, grouped as subclades of reference strain CP007462. All four WGS assemblies, and the MAG assemblies from samples S134 and S135 grouped together in a CP007462 subclade (Fig. 7; see Fig. S2A in the supplemental material). However, MAG assemblies, S169 and S170, had longer branch lengths than their corresponding WGS assemblies (Fig. 7; see Fig. S2A in the supplemental material). The WGS and MAG assemblies for seafood firm L S125 grouped as a subclade of reference genome CP015593; their branch lengths were considerably long due to low SNP coverage (Fig. 8; see Fig. S2A in the supplemental material).

**Figure 8.**
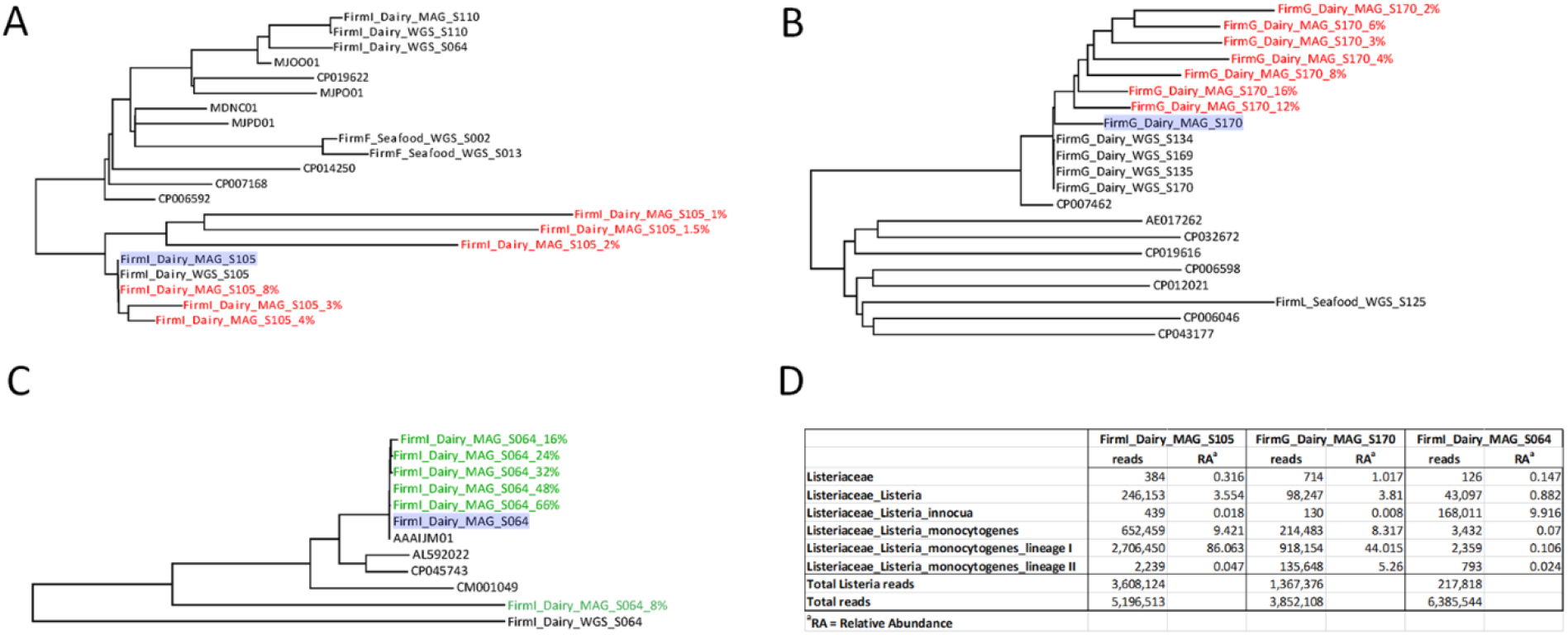
In silico fastq dilutions of 3 metagenomic assembled *Listeria* genomes (A, B, C). The first sample, FirmI_Dairy_MAG_S105 (A) had both a high relative abundance of *L. monocytogenes* and a significant number of *L. monocytogenes* lineage I reads. The second sample, FirmG_Dairy_MAG_S170 (B) had a high relative abundance and number of *L. monocytogenes* lineage I reads but also exhibited reads for *L. monocytogenes* lineage II. The third and final sample, FirmI_Dairy_MAG_S064 (C) was culture positive for both *L. moncytogenes* and *L. innocua* but shotgun metagenomic sequencing and analysis only identified reads and relative abundance for *L. innocua*. Panel D indicates the sequencing reads and relative abundance data mapped to *Listeria* for all 3 samples.

The lineage II *L*. *monocytogenes* SNP clade is comprised of 13 matching WGS and MAG assemblies within four SNP subclades (Fig. 7; see Fig. S2B in the supplemental material). Three samples S014, S073, and S077, collected from seafood firm F produced WGS and corresponding MAG assemblies that grouped with reference strain CP020828 (Fig. 7; see Fig. S2B in the supplemental material). Four samples from seafood firm A grouped with CP020828. MAG S037 from seafood firm A did not group with the corresponding lineage II WGS S037 assembly (Fig. 7; see Table S4 in the supplemental material). The other three MAG assemblies from seafood firm A, S072, S096 and S099, grouped with their corresponding WGS assemblies, as well as the assemblies from firm F in the CP020828 subclade (Fig. 7; see Fig. S2B in the supplemental material).

The lineage II SNP clade also has two subclades with samples collected from dairy firms J and G. The WGS and MAG assemblies from all the samples from these firms grouped in a subclade with reference strain CP007687 (Fig. 7, see Fig. S2B in the supplemental material). The second dairy firm subclade is comprised of reference strain CP007199 and the WGS assembly of firm G, S051 (Fig. 7, see Fig. S2B in the supplemental material). MAG S051 could not be placed in the SNP tree due to low genome coverage (Fig. 7, see Table S3 and Table S4 in the supplemental material).

The final lineage II subclade, with reference strain CP025560, is comprised of five samples from seafood firm L, and one from seafood firm W (Fig. 7; see Fig. S2B in the supplemental material). Three samples from seafood firm L, S108, S123, and S097 produced WGS and MAG assemblies that grouped together in this subclade (Fig. 7; Fig. S2B in the supplemental material). Seafood firm L MAG assembly S032 placed outside of the CP025560 subclade, while the corresponding WGS assembly is grouped within (Fig. 7; see Fig. S2B in the supplemental material). SNP analysis of the MAG S123, which was culture negative for *L*. *monocytogenes*, assembled and grouped within the CP025560 lineage II subclade (Fig. 7; see Fig. S2B in the supplemental material). The final sample in the CP025560 subclade is Sample S082from seafood firm W, which produced WGS and MAG assemblies that grouped in the subclade (Fig. 7; see Fig. S2B in the supplemental material)

### *In silico* Analysis of Shotgun Metagenomes

The comparison of lineage I and II *L*. *monocytogenes* WGS isolate genomes to corresponding MAG assemblies from parallel quasi-metagenomes revealed disparities in SNP phylogenies of some MAG assemblies (Fig. 7; see Fig. S2 in the supplemental material). In some samples, insufficient genome coverage precluded the placement of a MAG assembly into the SNP tree. Having the WGS assemblies from pure isolate genomes allowed us to predict the lineage of the *L*. *monocytogenes* in each culture enrichment bacterial community. However, the assembly of bacterial genomes from a metagenome is confounded by the presence of sequence reads from other bacterial genomes, especially those that are present in higher abundances. Additionally, metagenomic datasets may have multiple lineages and species of some genera, such as those that are closely related, that reduce the number of unique SNPs, resulting in longer branch lengths due to inadequate genome coverage. Here we performed an *in silico* downsampling study on three swab enrichment sample metagenomes to improve our understanding of how reducing the number of lineage specific *Listeria* sequence reads in a culture positive metagenomic enrichment effects SNP phylogeny.

The first metagenomic dataset we analyzed, MAG S105 from dairy firm I, was collected from a zone 3 floor drain in room 1 of the firm lineage I *L*. *monocytogenes* was cultured from the UVM enrichment. MAG S105 contained 2,706,450 *L. monocytogenes* sequence reads (86% proportional abundance), which were sequentially downsampled *in silico* to create seven new sequence read datasets using from 0.5% to 8% of the original read set (Fig. 8A and 8D). The S105 dataset with 0.5% of sequence reads was missing all lineage I *L*. *monocytogenes* SNPs and could not be placed into the S105 SNP tree (Fig. 8A; see Table S5 in the supplemental material). Datasets downsampled to 1%, 1.5%, and 2% of starting sequence reads were missing 12,981, 5,381 and 1,471 *L. monocytogenes* SNPs, respectively resulting in genome coverages of 3.4X (0.5%), 3.8X (1.5%) and 4.4X (2%). These three *in silico* downsampled datasets resulted in correct placement in the MAG S105 SNP cluster but had long branch lengths, in this case due to reduced genome coverage and missing SNPs (Fig. 8A; see Table S5 in the supplemental material). The S105 MAGs with sequence reads downsampled to 3% and 4%, equivalent to genome coverages of 6.4X and 8.4X respectively, were placed in a polyphyletic SNP cluster with the original MAG and corresponding WGS S105 genome assembly (Fig. 8A; see Table S5 in the supplemental material) Finally, the read dataset downsampled to 8%, which had zero missing SNPs, grouped with the original MAG and the corresponding WGS S105 genome assembly from firm I. The genome coverage for the 8% and original sequence read files were 16.9X (290,711 *L. monocytogenes* sequence reads) and 248X (3,623,130 *L. monocytogenes* sequence reads) respectively (Fig. 8A; see Table S5 in the supplemental material).

The second *in silico* reduction series was performed with dairy firm G MAG S170 collected in zone 3 on the top of a stair leading to a cheese cave. Dairy firm G MAG S170 had a 44% proportional abundance of lineage I *L. monocytogenes*, equivalent to 918,154 sequence reads, and a 5.3% proportional abundance of lineage II *L. monocytogenes* (135,648 sequence reads) (Fig. 8D; see Table S3 in the supplemental material). *In silico* downsampling of MAG S170 generated sequence datasets with 2%, 3%, 4%, 6%, 8%, 12% and 16% of total reads. The genome coverage of the original MAG S170 assembly was low (21X with 3,463 missing SNPs) compared to the original dairy firm I MAG assembly S105 (248X with 0 missing SNPs) (see Table S5 in the supplemental material). This low genome coverage resulted in a MAG S170 subclade of the corresponding lineage I WGS S170 assembly, and three additional lineage I WGS assemblies from Firm G (Fig. 8B; see Table S5 in the supplemental material). Furthermore, the reduced MAG S170 grouped with the original MAG S170 assembly, with branch lengths that increased as sequence reads were reduced (Fig. 8B). The low genome coverage in MAG S170 is likely due to the presence of lineage II sequence reads, which impact SNP clustering. The additional WGS assemblies, S134, S135 and S169 that grouped with WGS S170 were from swabs collected from the stairs and floor in zone 3 of the cheese cave area in firm G (Fig. 8B; see Table S5 in the supplemental material).

The third set of reduced read MAG assemblies were from another firm I sample, S064, collected from the floor under a piece of equipment in room 2 of the firm. The environmental swab culture enrichment for MAG S064 was culture positive for lineage I *L. moncytogenes* and *L. innocua*. Shotgun metagenomic sequencing produced a microbiome with proportional abundances of 10% *L*. *innocua* and <1% lineage I *L*. *monocytogenes*, equating to 168,011 and 2,359 sequencing reads, respectively (Fig. 8D). *In silico* reduction of MAG S064 produced read datasets with 66%, 48%, 32%, 24% 16% and 8% total reads, and all except the 8% dataset grouped in a SNP cluster with the original MAG S064 sample and AAAIJM01, an *L. innocua* reference isolate; the *L*. *innocua* isolate from S064 was not sequenced for the GenomeTrakr program (Fig. 8C). *L*. *innocua* genome coverages for the fastq files of MAG S064 were 22X (66% fastq), 17X (48% fastq), 13X (32% fastq), 11X (24% fastq), 10X (16% fastq), and 8X (8% fastq); coverage for the original fastq file was 31X (Sup Table S4). Both the 8% MAG S064 with 18,853 *Listeria* reads and 275,743 *L*. *innocua* missing SNPs, and the *L*. *monocytogenes* WGS isolate from sample S064, grouped outside of the *L*. *innocua* SNP cluster (Fig. 8C and 8D, see Table S5 in the supplemental material).

## DISCUSSION

Metagenomics is a powerful tool for characterizing microbial communities and the translation of ‘omics’ technologies like this to food microbiology will have a significant impact in the food industry and for public health (29, 30). The applications of this technology extend far beyond just public health, they can also provide valuable insights about food quality, and there is evidence that the microbiome is likely an important and effective hazard indicator within the food supply chain (31).

We expected to discover significant differences between the enriched microbiomes associated with dairy and seafood food production facilities, due to the different types of commodities produced in each type of facility. In other studies, it has been reported that food ingredients can act as seeding sources for members of the food production environment microbiome (2, 32). We did not observe any significant differences when looking at the alpha diversity among swabs taken from each firm, even though we observed variability between firms and between individual swabs. This might be largely due to the sample custody requirement that did not permit access to culture independent microbiomes in these facilities. It is well documented that culture enrichment tends to reduce diversity of sample populations as culture conditions, even in non-selective media, tend to favor those organisms that are not fastidious (16, 22, 33, 34). The culture enrichment conditions used in this study, although selective for *Listeria monocytogenes*, allowed growth of selected other organisms. It is interesting to note that we did observe increased diversity in swab enrichment microbiomes when *L*. *monocytogenes* was present. In many, the relative abundance of *L. monocytogenes* was quite low (< 10%) due to increased growth of indigenous bacteria. This contrasts with the more favorable outcome when the selective supplements in UVM favor *Listeria* growth and thus lower alpha diversity.

*L. monocytogenes* is a frequently isolated and sometimes a persistent contaminant within food industry settings (35–37). As such, extensive efforts are made to control this pathogen throughout the food supply chain, especially in food production environments, where *Listeria* is commonly transferred to food products by cross contamination from surfaces in production facilities (38, 39). Additionally, the sanitization and disinfection methods employed to prevent cross contamination can provide niches and possibly influence the formation of biofilm communities that may enhance bacterial survival in built food production facilities (37, 39, 40). *Pseudomonas* species were highly abundant in our environmental swab culture enrichments from both dairy and seafood production facilities (Fig. 4, Fig.5, see Table S2, S3 in the supplemental material). While it was not unusual to see high proportional abundances of *Pseudomonas* in foods or food environments, studies have shown that *Pseudomonas* species and *L. monocytogenes* can co-exist in mixed-species biofilms (41–45). Additional sampling studies are needed in food production environments to demonstrate that co-existence and mixed species biofilms contribute to food and food facility contamination and harborage. Our observation of the identical lineage II *L*. *monocytogenes* genomes in MAGs and WGS sequences generated from samples collected 8 months apart in seafood firm L, suggests points of harborage in this firm (Fig. 7; see Fig. S2B in the supplemental material). Details about sanitation and disinfection schedules during this time could provide additional data to understand the effects of sanitizers and disinfectants on harborage and persistence facility microbiotas over time.

The dominant taxa observed across all samples were *Enterococcus*, *Pseudomonas*, *Lactococcus*, and *Listeria*. Additionally, *Brochothrix* and *Carnobacterium* were present in swab enrichment samples from seafood facilities as dominant enriched microbiome members. There was a high level of variability in relative abundance from swab-to-swab enrichment microbiomes, even for these dominant taxa, and it was not possible to attribute a certain community or taxa with a facility type, specific zone or sample location, with statistical confidence.

In addition to the dominant taxa listed above, targeted amplicon sequencing also revealed taxa in the lower abundant phyla, such as Fusobacteria, Actinobacteria, and Bacteroidetes, that were uniquely enriched in selective UVM cultures from some firms. For example, *Fusobacterium* was the only genus identified in the Fusobacteria Phylum. *Fusobacterium* is a commensal bacterium of clinical significance in humans and animals (see Table S2 in the supplemental material) (46–50). Our 16S rRNA amplicon data suggests that *F*. *ulcerans* is the species colonizing these firms, although we did not confirm this. The identification of *Fusobacterium* 16S rRNA amplicon sequences in 100% of samples from Firm O and 3% to 38% of samples from firms AM, G, I and F is unusual since this genus consists mostly of obligate anaerobes, except for *F*. *nucleatum*, which has been demonstrated to survive aerotolerant conditions (51, 52). This finding suggests that anaerobic niches may exist in areas that were sampled in these firms and the subsequent UVM culture enrichment provided a low oxygen environment allowing *Fusobacterium* to grow. *Fusobacterium* species are widely known for their pathogenic attributes in anaerobic biofilms and abscesses common to periodontal disease (*F*. *nuceatum*), lameness in hooved animals such as dairy cattle (*F*. *necrophorum*) and pathogenesis in Atlantic salmon (Fusobacteria sp.) (46–50, 52). The sources of Fusobacteria in these five firms are unknown. However, since dairy firms AM, G, and O manufacture cheese and firm F manufactures seafood, a possible source could be the ingredients originating from dairy cattle (the common one being milk) and fish that were brought into the firms. Further investigation is required to confirm this.

The proportional abundances of Bacteroidetes in the dairy and seafood swab samples were also low, only reaching 8%. *Dysgonomonas* species, which are known to contribute to polysaccharide utilization in human and insect guts, were the most prevalent Bacteroidetes genus (53, 54). Firm O had the highest number of swab samples with *Dysgonomonas* 16S rRNA sequence reads. Since most of these samples were collected from the floors of the firms, it is possible that the presence of *Dysgonomonas* could indicate the presence of insects in these firms.

The significant differences between enriched bacterial communities in the cleaning and processing rooms of dairy firm AM were likely due to differences in *Lactococcus* proportional abundances, which were higher in area G (33%), compared to areas A (0.09%), B (0.01%), C (0.02%) and D (4%). This is not surprising since area G samples were collected from zones 1 (food contact surfaces) or 2 (the handles and outside of mixing bowls, and commonly touched areas of mixers) where high abundances of *Lactococcus* would be expected in a cheese manufacturing facility. None of the swab samples in area G of firm AM were culture positive for *L. monocytogenes*, and most of the culture positive swab enrichments (18/21) were collected from zone 3 surfaces such as floors, floor drains or wheels of moving equipment or carts. Perhaps our 16S rRNA sequencing of bacterial community fingerprints in different areas can support efforts to track harborage and transfer patterns of *L. monocytogenes* in this and other production facilities.

The *L*. *monocytogenes* isolate WGS genome assemblies allowed us to assess the accuracy of SNP clustering of MAGs assembled from the same parent culture enrichments. We also performed *in silico* downsampling of datasets with varied proportional abundances of *L*. *monocytogenes* allowing us to begin to establish some limits for *Listeria* detection and subtyping using shotgun metagenomics. Pathogen detection was achieved for all downsampled datasets, even those with only 3x genome coverage. Not surprisingly, subtyping resolution increased as the number of *Listeria* reads increased. To summarize, a lineage I subcluster from dairy firm AM, had MAG assemblies with 1.1% (S011) and 0.8% (S002) proportional abundances of *L*. *monocytogenes* that grouped outside of their subclades. However, the corresponding WGS assemblies were grouped within the subclade; samples within the subclade had proportional abundances of *L*. *monocytogenes* ranging from 2% to 64%. In general, we observed good subclade assignment with as low as 6x genome coverage, although this was impacted by the presence of a second *L*. *monocytogenes* strain or *L*. *innocua*.

As the technology for next generation sequencing continues to advance, metagenomics will become a revolutionary tool for improving food safety in regulatory laboratories. In the very near future, metagenomics will become a reliable screening tool for analyzing the entire microbial community in environmental swabs and food matrices. Long read sequencing technologies applied to culture independent metagenomic sequencing will usher in a new era of screening and detecting foodborne pathogens, providing a complete analysis of microbial communities. This study provides evidence that routine analyses with metagenomic methods in food testing laboratories could enhance foodborne outbreak prevention and the resolution of ongoing outbreaks.

## ACKNOWLEDGMENTS

We would like to thank the FDA Northeast Food and Feed Laboratory for providing the samples we used in this study. We thank the FDA’s Office of the Chief Scientist for funding this work as part of the project “MetagenomeTrakr pilot program for rapid foodborne pathogen detection”. Brandon Kocurek was an Oak Ridge Institute for Science and Education fellow, and we also thank the Department of Energy for their support. The opinions expressed in this paper are those of the authors and not necessarily those of the Food and Drug Administration. Use of trade names is for identification only and does not imply endorsement by the U. S. Food and Drug Administration, or by the Public Health Service, or by the U.S. Department of Health and Human Services.

## REFERENCES

1. De Filippis F, Valentino V, Alvarez-Ordóñez A, Cotter PD, Ercolini D. 2021. Environmental microbiome mapping as a strategy to improve quality and safety in the food industry. Current Opinion in Food Science 38:168–176.

2. Cobo-Díaz JF, Alvarez-Molina A, Alexa EA, Walsh CJ, Mencía-Ares O, Puente-Gómez P, Likotrafiti E, Fernández-Gómez P, Prieto B, Crispie F, Ruiz L, González-Raurich M, López M, Prieto M, Cotter P, Alvarez-Ordóñez A. 2021. Microbial colonization and resistome dynamics in food processing environments of a newly opened pork cutting industry during 1.5 years of activity. Microbiome 9:204.

3. Møretrø T, Langsrud S. 2017. Residential Bacteria on Surfaces in the Food Industry and Their Implications for Food Safety and Quality. Compr Rev Food Sci Food Saf 16:1022–1041.

4. Konya T, Scott JA. 2014. Recent Advances in the Microbiology of the Built Environment. Current Sustainable/Renewable Energy Reports 1:35–42.

5. Wang Y, Pettengill JB, Pightling A, Timme R, Allard M, Strain E, Rand H. 2018. Genetic Diversity of Salmonella and Listeria Isolates from Food Facilities. Journal of Food Protection 81:2082–2089.

6. Allard MW, Strain E, Rand H, Melka D, Correll WA, Hintz L, Stevens E, Timme R, Lomonaco S, Chen Y, Musser SM, Brown EW. 2019. Whole genome sequencing uses for foodborne contamination and compliance: Discovery of an emerging contamination event in an ice cream facility using whole genome sequencing. Infection, Genetics and Evolution 73:214–220.

7. Timme RE, Sanchez Leon M, Allard MW. 2019. Utilizing the Public GenomeTrakr Database for Foodborne Pathogen Traceback. Methods Mol Biol 1918:201–212.

8. De Filippis F, La Storia A, Villani F, Ercolini D. 2013. Exploring the sources of bacterial spoilers in beefsteaks by culture-independent high-throughput sequencing. PLoS One 8:e70222.

9. Stellato G, De Filippis F, La Storia A, Ercolini D. 2015. Coexistence of Lactic Acid Bacteria and Potential Spoilage Microbiota in a Dairy Processing Environment. Appl Environ Microbiol 81:7893–904.

10. Bokulich NA, Ohta M, Richardson PM, Mills DA. 2013. Monitoring Seasonal Changes in Winery-Resident Microbiota. PLoS One 8:e66437.

11. Bokulich NA, Bergsveinson J, Ziola B, Mills DA. 2015. Mapping microbial ecosystems and spoilage-gene flow in breweries highlights patterns of contamination and resistance. eLife 4:e04634.

12. Dzieciol M, Schornsteiner E, Muhterem-Uyar M, Stessl B, Wagner M, Schmitz-Esser S. 2016. Bacterial diversity of floor drain biofilms and drain waters in a Listeria monocytogenes contaminated food processing environment. International Journal of Food Microbiology 223:33–40.

13. Møretrø T, Moen B, Heir E, Hansen A, Langsrud S. 2016. Contamination of salmon fillets and processing plants with spoilage bacteria. Int J Food Microbiol 237:98–108.

14. Flores GE, Bates ST, Caporaso JG, Lauber CL, Leff JW, Knight R, Fierer N. 2013. Diversity, distribution and sources of bacteria in residential kitchens. Environ Microbiol 15:588–96.

15. Buytaers FE, Saltykova A, Denayer S, Verhaegen B, Vanneste K, Roosens NHC, Piérard D, Marchal K, De Keersmaecker SCJ. 2020. A Practical Method to Implement Strain-Level Metagenomics-Based Foodborne Outbreak Investigation and Source Tracking in Routine. Microorganisms 8:1191.

16. Ottesen A, Ramachandran P, Reed E, White JR, Hasan N, Subramanian P, Ryan G, Jarvis K, Grim C, Daquiqan N, Hanes D, Allard M, Colwell R, Brown E, Chen Y. 2016. Enrichment dynamics of Listeria monocytogenes and the associated microbiome from naturally contaminated ice cream linked to a listeriosis outbreak. BMC Microbiol 16:275.

17. Leonard SR, Mammel MK, Lacher DW, Elkins CA. 2016. Strain-Level Discrimination of Shiga Toxin-Producing Escherichia coli in Spinach Using Metagenomic Sequencing. PLoS One 11:e0167870.

18. Chen LX, Anantharaman K, Shaiber A, Eren AM, Banfield JF. 2020. Accurate and complete genomes from metagenomes. Genome Res 30:315–333.

19. Meziti A, Rodriguez-R LM, Hatt JK, Peña-Gonzalez A, Levy K, Konstantinidis KT, McBain AJ. 2021. The Reliability of Metagenome-Assembled Genomes (MAGs) in Representing Natural Populations: Insights from Comparing MAGs against Isolate Genomes Derived from the Same Fecal Sample. Applied and Environmental Microbiology 87:e02593–20.

20. Hyeon JY, Li S, Mann DA, Zhang S, Li Z, Chen Y, Deng X. 2018. Quasimetagenomics-Based and Real-Time-Sequencing-Aided Detection and Subtyping of Salmonella enterica from Food Samples. Appl Environ Microbiol 84.

21. Hyeon JY, Mann DA, Townsend AM, Deng X. 2018. Quasi-metagenomic Analysis of Salmonella from Food and Environmental Samples. J Vis Exp doi:10.3791/58612.

22. Ottesen A, Ramachandran P, Chen Y, Brown E, Reed E, Strain E. 2020. Quasimetagenomic source tracking of Listeria monocytogenes from naturally contaminated ice cream. BMC Infect Dis 20:83.

23. Townsend A, Li S, Mann DA, Deng X. 2020. A quasimetagenomics method for concerted detection and subtyping of Salmonella enterica and E. coli O157:H7 from romaine lettuce. Food Microbiol 92:103575.

24. Wagner E, Fagerlund A, Langsrud S, Møretrø T, Jensen MR, Moen B. 2021. Surveillance of Listeria monocytogenes: Early Detection, Population Dynamics, and Quasimetagenomic Sequencing during Selective Enrichment. Appl Environ Microbiol 87:e0177421.

25. Magoc T, Salzberg SL. 2011. FLASH: fast length adjustment of short reads to improve genome assemblies. Bioinformatics 27:2957–63.

26. Edgar RC. 2010. Search and clustering orders of magnitude faster than BLAST. Bioinformatics 26:2460–1.

27. Matias Rodrigues JF, Schmidt TSB, Tackmann J, von Mering C. 2017. MAPseq: highly efficient k-mer search with confidence estimates, for rRNA sequence analysis. Bioinformatics 33:3808–3810.

28. Kumar S, Tamura K, Nei M. 2004. MEGA3: Integrated software for Molecular Evolutionary Genetics Analysis and sequence alignment. Brief Bioinform 5:150–63.

29. Brown EW, Gonzalez-Escalona N, Stones R, Timme R, Allard MW. 2017. The Rise of Genomics and the Promise of Whole Genome Sequencing for Understanding Microbial Foodborne Pathogens, p 333-351. *In* Gurtler JB, Doyle MP, Kornacki JL (ed), Foodborne Pathogens: Virulence Factors and Host Susceptibility doi:10.1007/978-3-319-56836-2_11. Springer International Publishing, Cham.

30. Walsh AM, Crispie F, Claesson MJ, Cotter PD. 2017. Translating Omics to Food Microbiology. Annu Rev Food Sci Technol 8:113–134.

31. Beck KL, Haiminen N, Chambliss D, Edlund S, Kunitomi M, Huang BC, Kong N, Ganesan B, Baker R, Markwell P, Kawas B, Davis M, Prill RJ, Krishnareddy H, Seabolt E, Marlowe CH, Pierre S, Quintanar A, Parida L, Dubois G, Kaufman J, Weimer BC. 2020. Monitoring the microbiome for food safety and quality using deep shotgun sequencing. bioRxiv doi:10.1101/2020.05.18.102574:2020.05.18.102574.

32. Johnson J, Curtin C, Waite-Cusic J. 2021. The Cheese Production Facility Microbiome Exhibits Temporal and Spatial Variability. Frontiers in Microbiology 12.

33. Jarvis KG, White JR, Grim CJ, Ewing L, Ottesen AR, Beaubrun JJ, Pettengill JB, Brown E, Hanes DE. 2015. Cilantro microbiome before and after nonselective pre-enrichment for Salmonella using 16S rRNA and metagenomic sequencing. BMC Microbiol 15:160.

34. Jarvis KG, Hsu CK, Pettengill JB, Ihrie J, Karathia H, Hasan NA, Grim CJ. 2021. Microbiome Population Dynamics of Cold-Smoked Sockeye Salmon during Refrigerated Storage and after Culture Enrichment. J Food Prot 85:238–253.

35. Osek J, Lachtara B, Wieczorek K. 2022. Listeria monocytogenes – How This Pathogen Survives in Food-Production Environments? Frontiers in Microbiology 13.

36. Malley TJV, Butts J, Wiedmann M. 2015. Seek and Destroy Process: Listeria monocytogenes Process Controls in the Ready-to-Eat Meat and Poultry Industry. Journal of Food Protection 78:436–445.

37. Tan X, Chung T, Chen Y, Macarisin D, LaBorde L, Kovac J. 2019. The occurrence of Listeria monocytogenes is associated with built environment microbiota in three tree fruit processing facilities. Microbiome 7:115.

38. Ferreira V, Wiedmann M, Teixeira P, Stasiewicz MJ. 2014. Listeria monocytogenes persistence in food-associated environments: epidemiology, strain characteristics, and implications for public health. J Food Prot 77:150–70.

39. Giaouris E, Heir E, Hébraud M, Chorianopoulos N, Langsrud S, Møretrø T, Habimana O, Desvaux M, Renier S, Nychas GJ. 2014. Attachment and biofilm formation by foodborne bacteria in meat processing environments: causes, implications, role of bacterial interactions and control by alternative novel methods. Meat Sci 97:298–309.

40. Fagerlund A, Langsrud S, Møretrø T. 2021. Microbial diversity and ecology of biofilms in food industry environments associated with Listeria monocytogenes persistence. Current Opinion in Food Science 37:171–178.

41. Heir E, Møretrø T, Simensen A, Langsrud S. 2018. Listeria monocytogenes strains show large variations in competitive growth in mixed culture biofilms and suspensions with bacteria from food processing environments. Int J Food Microbiol 275:46–55.

42. Pang X, Yuk HG. 2019. Effects of the colonization sequence of Listeria monocytogenes and Pseudomonas fluorescens on survival of biofilm cells under food-related stresses and transfer to salmon. Food Microbiol 82:142–150.

43. Puga CH, Dahdouh E, SanJose C, Orgaz B. 2018. Listeria monocytogenes Colonizes Pseudomonas fluorescens Biofilms and Induces Matrix Over-Production. Frontiers in Microbiology 9.

44. Papaioannou E, Giaouris ED, Berillis P, Boziaris IS. 2018. Dynamics of biofilm formation by Listeria monocytogenes on stainless steel under mono-species and mixed-culture simulated fish processing conditions and chemical disinfection challenges. Int J Food Microbiol 267:9–19.

45. Fagerlund A, Møretrø T, Heir E, Briandet R, Langsrud S. 2017. Cleaning and Disinfection of Biofilms Composed of Listeria monocytogenes and Background Microbiota from Meat Processing Surfaces. Appl Environ Microbiol 83.

46. Austin B. 2011. Taxonomy of bacterial fish pathogens. Veterinary Research 42:20.

47. Maher M, Palmer R, Gannon F, Smith T. 1995. Relationship of a Novel Bacterial Fish Pathogen to Streptobacillus moniliformis and the Fusobacteria Group, based on 16S Ribosomal RNA Analysis. Systematic and Applied Microbiology 18:79–84.

48. Van Metre DC. 2017. Pathogenesis and Treatment of Bovine Foot Rot. Vet Clin North Am Food Anim Pract 33:183–194.

49. Bennett G, Hickford J, Zhou H, Laporte J, Gibbs J. 2009. Detection of Fusobacterium necrophorum and Dichelobacter nodosus in lame cattle on dairy farms in New Zealand. Res Vet Sci 87:413–5.

50. Han YW. 2015. Fusobacterium nucleatum: a commensal-turned pathogen. Current Opinion in Microbiology 23:141–147.

51. Brook I. 2017. 184 - Anaerobic Bacteria, p 1628-1644.e2. In Cohen J, Powderly WG, Opal SM (ed), Infectious Diseases (Fourth Edition) doi:https://doi.org/10.1016/B978-0-7020-6285-8.00184–2. Elsevier.

52. Diaz PI, Zilm PS, Rogers AH. 2002. Fusobacterium nucleatum supports the growth of Porphyromonas gingivalis in oxygenated and carbon-dioxide-depleted environments. Microbiology 148:467–472.

53. Kmezik C, Mazurkewich S, Meents T, McKee LS, Idström A, Armeni M, Savolainen O, Brändén G, Larsbrink J. 2021. A polysaccharide utilization locus from the gut bacterium Dysgonomonas mossii encodes functionally distinct carbohydrate esterases. J Biol Chem 296:100500.

54. Kucuk RA. 2020. Gut Bacteria in the Holometabola: A Review of Obligate and Facultative Symbionts. Journal of Insect Science 20.

